# Daylong Acoustic Recordings of Grazing and Rumination Activities in Dairy Cows

**DOI:** 10.1101/2023.10.18.562979

**Authors:** Luciano S. Martinez-Rau, José O. Chelotti, Mariano Ferrero, Santiago A. Utsumi, Alejandra M. Planisich, Leandro D. Vignolo, Leonardo L. Giovanini, H. Leonardo Rufiner, Julio R. Galli

**Affiliations:** Instituto de Investigación en Señales, Sistemas e Inteligencia Computacional, sinc(i), FICH-UNL/CONICET, Argentina; Department of Computer and Electrical Engineering, Mid Sweden University, Sundsvall, Sweden; TERRA Teaching and Research Center, University of Liège, Gembloux Agro-Bio Tech (ULiège-GxABT), 5030 Gembloux, Belgium; W.K. Kellogg Biological Station and Department of Animal Science, Michigan State University, United States; Department of Animal and Range Science, New Mexico State University, United States; Facultad de Ciencias Agrarias, Universidad Nacional de Rosario, Argentina; Laboratorio de Cibernética, Facultad de Ingeniería, Universidad Nacional de Entre Ríos, Argentina; Instituto de Investigaciones en Ciencias Agropecuarias de Rosario, IICAR, UNR-CONICET, Argentina

## Abstract

Monitoring livestock feeding behavior may help assess animal welfare and nutritional status, and to optimize pasture management. The need for continuous and sustained monitoring requires the use of automatic techniques based on the acquisition and analysis of sensor data. This work describes an open dataset of acoustic recordings of the foraging behavior of dairy cows. The dataset includes 662 h of daily records obtained using unobtrusive and non-invasive instrumentation mounted on five lactating multiparous Holstein cows continuously monitored for six non-consecutive days in pasture and barn. Labeled recordings precisely delimiting grazing and rumination bouts are provided for a total of 400 h and for over 6,200 ingestive and rumination jaw movements. Companion information on the audio recording quality and expert-generated labels is also provided to facilitate data interpretation and analysis. This comprehensive dataset is a useful resource for studies aimed at exploring new tools and solutions for precision livestock farming.

## Background & Summary

Advances in information and communication technologies enabled the development of precision livestock farming (PLF) systems with potential applications to improve farm operational efficiency and animal welfare^1,2^. Over the last three decades, PLF has grown substantially, attracting farmers, operators and industries around the world^3,4^. New PLF developments include methodologies to enable the individual monitoring of livestock feeding behavior, which might be used to detect changes in animal welfare with direct insights into animal nutrition, health or performance^5–7^. Wearable sensors are the most common data acquisition method to monitor feeding behavior^8,9^. Accelerometers and inertial measurement units determine head and neck movements and have been used mainly in confined environments^10,11^. Acoustic sensors are typically preferred over motion sensors in free-ranging conditions^12^ to classify different animal jaw movements (JMs)^13–17^ and feeding behavior^18,19^. Furthermore, distinguishing different types of JMs is useful for delimiting grazing and rumination bouts^20^, estimating dry matter intake, and discriminating different feedstuffs and plants^21,22^.

The acoustic monitoring of foraging behavior is an engineering task that requires robust solutions capable of tolerating noise, interference and disturbance ^12^. The opportunities to use acoustic methods for practical farm-level management and animal research are ample^23^, but the limited availability of public/open acoustic datasets could hinder new and relevant research^24^. To the best of our knowledge, there are only two open datasets of cattle acoustic sounds. The first dataset contains 52 audio recordings of JMs of dairy cows grazing on two contrasting forage species at two sward heights^25^. The other dataset provides 270 samples of cattle calls, also called cattle vocalizations^26,27^.

This work presents a dataset of audio recordings of chewing and biting sounds of dairy cows along with their corresponding event identification labels. The dataset is organized into three groups. (*i*) It includes 24-h audio recordings of continuously monitored dairy cows grazing in pastures or visiting the dairy milking barn. A total of 662.5 h were recorded, from which 400.4 h corresponded to sounds registered in a free-range pasture environment. Annotations of the grazing and rumination bouts are provided for each of the cows. Periods during which the dairy cows were inside the dairy barn are also indicated. (*ii*) It contains two audio files of 54.6 min of grazing and 30.0 min of rumination, with the corresponding labels for JMs. Experts identified and labeled 4,221 ingestive JMs and 2,006 rumination JMs produced during grazing and rumination, respectively. (*iii*) It provides a comprehensive description of the different types of JMs and animal behaviors, and specific information about the audio recordings. The dataset presented here has been previously used to create automatic machine-learning algorithms for detecting and classifying different JMs^28–30^ and for classifying grazing and rumination activities^25,31–33^. This dataset could be used to improve the recognition rate, generalization ability, and noise robustness of existing algorithms^34^, as well as to develop novel algorithms that combine acoustic signals with other sources of information^35^.

## Methods

The field study took place from July 31 to August 19, 2014, and was conducted at the W. K. Kellogg Biological Station’s Pasture Dairy Research Center of Michigan State University, located in Hickory Corners, Michigan, US (GPS coordinates 42° 24^′^ 21.8^′′^ N 85° 24^′^ 08.4^′′^ W). The procedures for animal handling and care were revised and approved by the Institutional Animal Care and Use Committee of Michigan State University (#02*/*17 − 020 − 00) before the start of the experiment. As described by Watt et al.^36^, animals were managed on a grazing-based platform with free access to the robotic milking system. Voluntary milking (3.0 ± 1.0 daily milkings) was conducted using two Lely A3-Robotic milking units (Lely Industries NV, Maassluis, The Netherlands). Permissions for milking were set by a minimum expected milk yield of 9.1 kg or a 6 h milking interval. Thus, milking frequency varied across cows according to milk yield. Dairy cows were fed a grain-based concentrate at 1 to 6 kg per kg of extracted milk (daily maximum 12 kg/cow) during milking and through automatic feeders located inside the dairy milking barn. The neutral detergent fiber (NDF), net energy for lactation (NEL), and average crude protein (CP) of the grain-based concentrate pellet supplied (Cargill Inc, Big Lake, MN) were 2.05 Mcal/kg dry matter (DM), 99.4 g/kg DM, and, 193.0 g/kg DM respectively. Cows were allowed 24-h access to grazing paddocks with a predominance of orchardgrass (Dactylis glomerata), tall fescue (*Lolium arundinacea*) and white clover (*Trifolium repens*), or perennial ryegrass (*Lolium perenne*) and white clover. Two allocations of ∼ 15 kg/cow of fresh pasture were offered daily, from 10:00 to 22:00 and from 22:00 to 10:00 (GMT-5), resulting in an average daily offer of ∼ 30 kg of DM/cow. Allocations of fresh ungrazed pasture were made available at opposite sides of the farm (south and north) to entice cow traffic through the milking shed. Thirty readings of sward height (SH, *x*) along each paddock were conducted by a plate meter to estimate pre-grazing and post-grazing herbage biomass to ground level (*Y,Y* = 125*x*; *r*^2^ = 0.96). This equation was also developed and verified for similar swards. Across the 16 paddocks used in this study, the average pre-grazing herbage biomass was 2387 ± 302 kg DM/ha (19.2 ± 2.5 cm SH) and the average post-grazing herbage biomass was 1396 ± 281 kg DM/ha (11.2 ± 2.2 cm SH). Composite hand-plucked samples from the 16 paddocks were used to determine the 48 h in vitro digestibility of DM (IVDMD) (Daisy II, Ankom Technology Corp.), the acid (ADF) and neutral detergent fiber (NDF) (Fiber Analyzer, Ankom Technology Corp., Fairport, NY), the crude protein (CP) (4010 CN Combustion, Costech Analytical Technologies Inc., Valencia, CA), and the acid detergent lignin (ADL) content of consumed forages. The values of DM expressed in terms of g/kg for IVDMD, CP, NDF, ADF and ADL were 781 ± 30, 257 ± 20, 493 ± 45, 187 ± 25, 33 ± 8, respectively.

For this study, 5 lactating high-producing multiparous Holstein cows were selected from a herd of 146 Holstein cows and used to non-invasively acquire and record acoustic signals over 24-h periods. Specific characteristics of individual cows are provided in Table 1. Individualized 24-h audio recordings were conducted on July 31, and August 4, 6, 11, 13 and 18, 2014, respectively. Recordings were obtained following a 5 x 5 Latin-square design (Table 2) using 5 independent monitoring systems (halters, microphones and recorders) that were rotated daily across the 5 cows and throughout 6 non-consecutive recording days. This design was decided to control for differences of sound data associated with a particular cow, recording systems or experiment day. On the first day, each recording system was randomly assigned to each cow. On the sixth day, the recording systems were reassigned to cows using the same order that was used on the first day. No training in the use of the recording systems was deemed necessary before study onset. Recording problems were encountered with the recording system number 2. On the first day, the recording trial had to be stopped a few hours before completion because the recording system was unfastened from the cow. This trial was considered valid and was not repeated. On the sixth day, the recording system failed to register any sound because the microphone connector was disconnected from the recorder. This trial was repeated on the next day (August 19) to complete the recordings of the sixth day. Changes in the order and completion of recording trials should be considered when designating trial days as a random variable in the experimental design. The weather conditions during the study were registered by the National Weather Service Station located at the Kellogg Biological Station (Table 3).

**Table 1.** Specific traits and description of the dairy cows used to acquire the audio recordings. The measurements were carried out on the first day of the experiment.

| | Cow 1<br>(ID: 2936) | Cow 2<br>(ID: 2909) | Cow 3<br>(ID: 2948) | Cow 4<br>(ID: 21036) | Cow 5<br>(ID: 2976) | Mean ± standard deviation |
| --- | --- | --- | --- | --- | --- | --- |
| Weight [kg] | 653 | 651 | 674 | 657 | 663 | 659.7 ± 9.4 |
| Lactation number | 3 | 3 | 2 | 2 | 3 | 2.6 ± 0.5 |
| Days in milk [d] | 130 | 125 | 68 | 141 | 62 | 105.2 ± 37.3 |
| Milk yield [kg/d] | 35.4 | 37.8 | 44.1 | 40.3 | 44.0 | 40.3 ± 3.8 |

**Table 2.** Latin-square design for recording systems, cows and days.

| Cow | Days (Date) |  |  |  |  |  |
| --- | --- | --- | --- | --- | --- | --- |
|  | 1 (Jul 31) | 2 (Aug 4) | 3 (Aug 6) | 4 (Aug 11) | 5 (Aug 13) | 6 (Aug 18-19) |
| 1 (ID: 2936) | 1 | 2 | 3 | 4 | 5 | 1 |
| 2 (ID: 2909) | 2 | 3 | 4 | 5 | 1 | 2* |
| 3 (ID: 2948) | 3 | 4 | 5 | 1 | 2 | 3 |
| 4 (ID: 21036) | 4 | 5 | 1 | 2 | 3 | 4 |
| 5 (ID: 2976) | 5 | 1 | 2 | 3 | 4 | 5 |

\* Audio recording repeated on August 19 due to problems associated with the microphone connector and recorder.

**Table 3.**
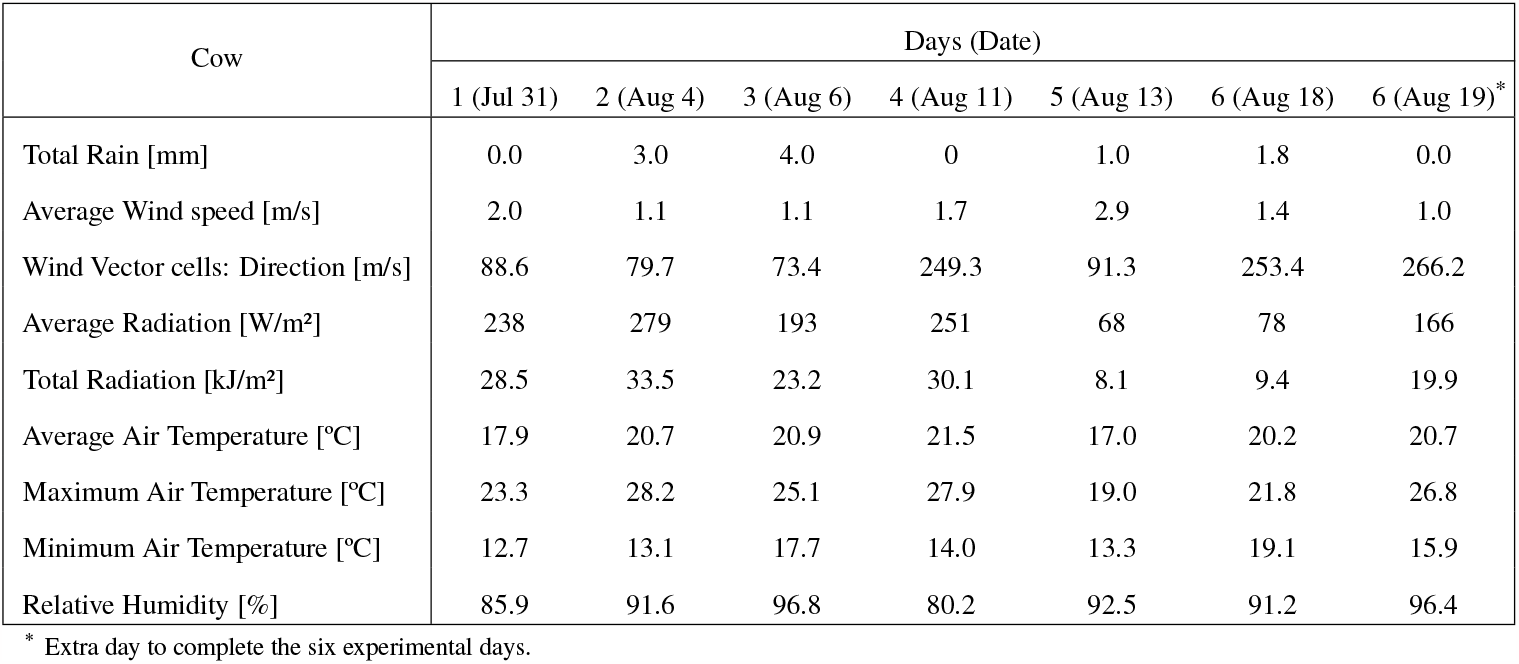
Weather conditions during audio recording trials.

| Cow | Days (Date) |  |  |  |  |  |  |
| --- | --- | --- | --- | --- | --- | --- | --- |
|  | 1 (Jul 31) | 2 (Aug 4) | 3 (Aug 6) | 4 (Aug 11) | 5 (Aug 13) | 6 (Aug 18) | 6 (Aug 19)* |
| Total Rain [mm] | 0.0 | 3.0 | 4.0 | 0 | 1.0 | 1.8 | 0.0 |
| Average Wind speed [m/s] | 2.0 | 1.1 | 1.1 | 1.7 | 2.9 | 1.4 | 1.0 |
| Wind Vector cells: Direction [m/s] | 88.6 | 79.7 | 73.4 | 249.3 | 91.3 | 253.4 | 266.2 |
| Average Radiation [W/m<sup>2</sup>] | 238 | 279 | 193 | 251 | 68 | 78 | 166 |
| Total Radiation [kJ/m<sup>2</sup>] | 28.5 | 33.5 | 23.2 | 30.1 | 8.1 | 9.4 | 19.9 |
| Average Air Temperature [°C] | 17.9 | 20.7 | 20.9 | 21.5 | 17.0 | 20.2 | 20.7 |
| Maximum Air Temperature [°C] | 23.3 | 28.2 | 25.1 | 27.9 | 19.0 | 21.8 | 26.8 |
| Minimum Air Temperature [°C] | 12.7 | 13.1 | 17.7 | 14.0 | 13.3 | 19.1 | 15.9 |
| Relative Humidity [%] | 85.9 | 91.6 | 96.8 | 80.2 | 92.5 | 91.2 | 96.4 |

\* Extra day to complete the six experimental days.

Each recording system consisted of two directional electret microphones connected to the stereo input channels of a digital recorder (Sony Digital ICD-PX312, Sony, San Diego, CA, USA). A 1.5 V AAA alkaline battery powered the digital recorder. The digital recorder saved the data in a 4 GB micro secure digital (SD) card (SanDisk SDSDB-004G-B35 SDHC, Western Digital, Milpitas, CA, USA). This instrumentation was enclosed in a weather proof protective case (1015 Micron Case Series, Pelican Products, Torrance, CA, USA) mounted to the top side of a halter neck strap (Fig. 1). One microphone was positioned facing inwards to capture the bone-transmitted vibrations and pressed against the forehead of the animal, while the other microphone faced outwards to capture the sounds produced by the animal. To achieve better microphone contact, hair of the central forehead area was removed using a sharp clipper. The microphones were held in the desired position by using a rubber foam and elastic headband attached to the halter. This design prevented microphone movements and allowed the insulation of microphones from environmental noise caused by wind, friction and scratches^37,38^.

**Figure 1.**
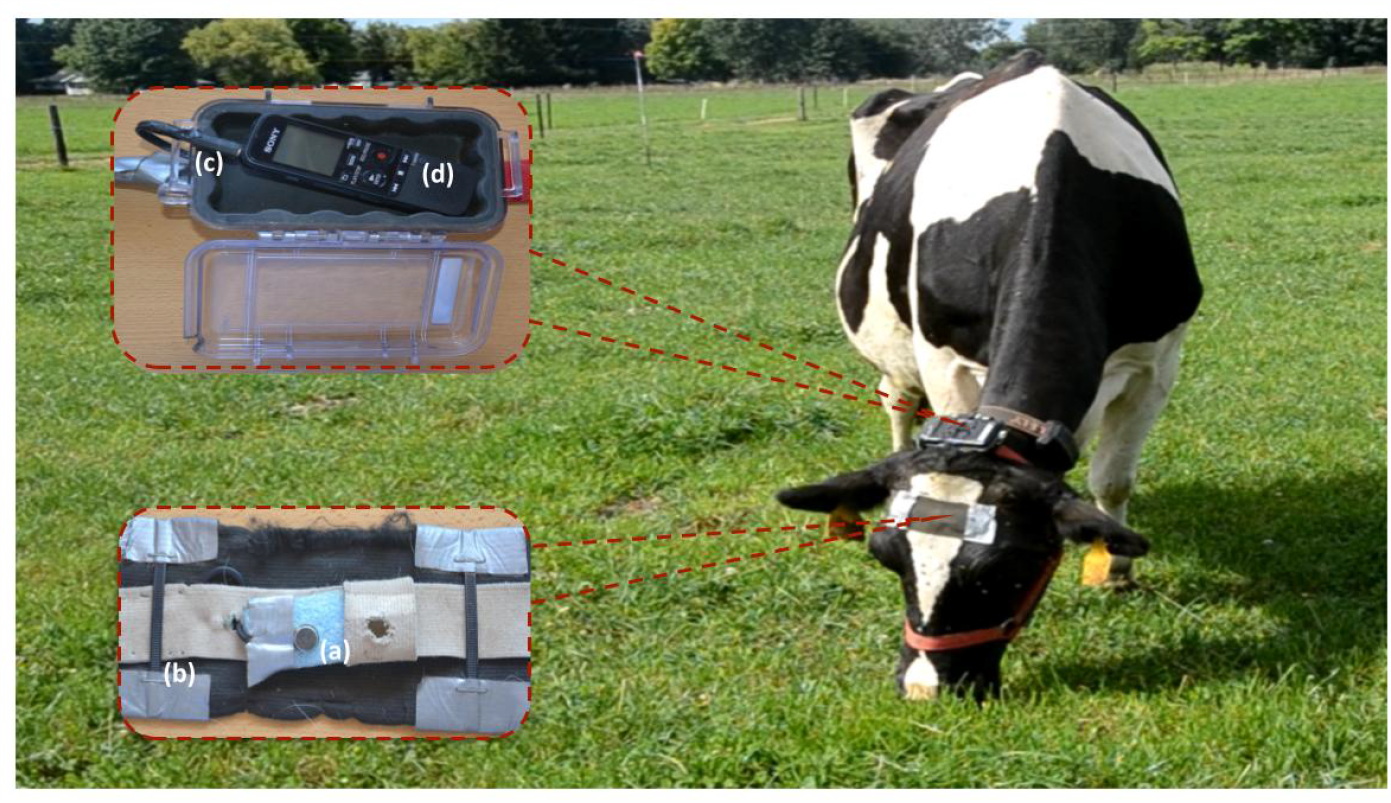
Recording system used to record the acoustic signals composed of inward and outward facing microphones (a). Wired microphones were covered by an elastic headband (b) and plugged (c) into a recorder housed inside a weather proof case attached to the top side of a halter neck strap (d).

After the morning milking session, the study cows were automatically separated into a holding pen. They were then restrained using head lockers to install recording systems equipped with new batteries and empty SD cards. As each cow completed the 24 h of continuous recording, they were manually guided to the head lockers to remove the recording systems. The date and relevant information of the recording systems and cows were kept in a logbook. A similar process was repeated on every trial day following the Latin-square design. In each recorder system, the two microphones were connected randomly to the stereo-input channels of the recorder at the beginning of trials. This information was not logged. Experienced animal handlers, who had extensive experience in animal behavior, data collection and analysis, directly observed the focal animals for blocks of ∼ 5 min each hour. Observation of foraging behavior, the time the equipment was turned on and other relevant parameters were documented and registered in the logbook. The handlers also checked the correct placement and location of recording systems on the cows. Observations were conducted at a distance from the animals to minimize disruptions of behavior.

The label files were generated by two experts with extensive experience in animal behavior understanding and digital analysis of audio signals^25,28,37–40^. The labeling was performed by an expert and the results were reviewed by another expert. The experts were guided by the logbook and used Audacity software^41^ to observe the sound waveforms and to listen to sounds to identify, classify, and label data into animal behavior categories. Annotations of interest that experts could not acoustically identify, such as the installation and removal of recording systems, were labeled by using the logbook registers. Although the experts matched all label assignments, there were some small differences in the start and/or end times (timestamp) of some labels. In those cases, both experts revised the labels together until they reached a mutual agreement. Additionally, as previously mentioned, the two microphones of each recording system were randomly connected to the stereo-input channels of the recorder throughout the trials. As a consequence, the stereo-input channels are swapped across the audio recordings. To address this, the experts listened to segments of grazing activity and barn location for all audio recordings and marked the one-to-one correspondence between the stereo-input channels and the two microphones (facing inwards and outwards of the forehead of the animal). However, establishing the proper microphone correspondence for some audio recordings was not straightforward due to the similar or wide variation in the channels. The experts made their decision based on a final mutual agreement.

Twenty-four-h recordings were registered in two settings: indoors, while cows visited the dairy milking barn and outdoors, while cows had free access to grazing pasture. During the continuous acoustic monitoring of cows, the animal handler annotated the rumination and feeding activities inside the milking barn in the logbook. However, the experts did not label these activities because the presence of acoustic noise in the audio recordings made it difficult to ensure their proper delimitation. The main focus of the experiment was to collect acoustic signals of foraging behavior while cows grazed in free-range conditions.

A total of 6,227 ingestive and rumination JMs were individualized, delimited and labeled by the experts, following the same approach and criteria used for labeling the animal behavior categories. This is a complex task that requires significant processing time and expertise in audio signal processing and inspection. Therefore, the start and end timestamps of the JMs could be subjective and may vary from the true bounds of the JMs in the audio files. To address this potential bias, an additional group of JMs’ timestamps was generated using a Python script. This script automatically adjusts the start and end boundaries of the JMs defined by the experts, without changing the JM label. Adjusted timestamps are determined based on the sound intensity during the JMs when it exceeds a threshold level. This threshold level is defined using the sound intensity during the pauses that occur between consecutive JMs.

Moreover, a pattern recognition JMs classifier algorithm^39^ has been used to automatically create JMs’ timestamps and labels in all grazing and rumination bouts of the daily recordings. The algorithm inputs were the channel corresponding to the facing-inward microphone of the audio recording and the outputs were a series of label files. The algorithm labels three types of JMs in terms of chews, bites and chew-bites. A post-processing algorithm was applied to have four types of JMs by dividing the chews into chews during grazing and chews during rumination. The JMs label files were not verified by the experts. Therefore, these files may contain possible identifications of JMs that did not exist, misidentifications of JMs that do exist, and/or incorrect JM labels.

## Data Records

The data is available at Figshare^42^. The audio recordings were saved in MPEG-1 Audio Layer III (MP3) format^43^ with a sampling rate of 44.1 kHz, providing a nominal recording bandwidth of 22 kHz and a dynamic range of 96 dB. The recordings were made in stereo, using one microphone per channel with a resolution of 16 bits at 192 kbps. This configuration made it possible to save up to 48 h of audio on the SD card with a battery autonomy of 55 h ensuring the desired 24-h recording with a good margin. The digital recorder automatically crops and generates a new MP3 file if the current audio recording is longer than 6 h. Thus, 24 h audio recordings are partitioned into 4 parts of approximately 6 h each.

The dataset is organized into three distinct groups (Fig. 2) as follows:

1. Daily recordings: It contains 30 ZIP files that correspond to the different recording trials of this study (6 days and 5 cows). Each ZIP file comprises ∼ 24 h of audio recordings and the corresponding activity label and automatically generated JM label files (Fig. 2). A total of 662.5 h are included in the 133 audio recordings, consisting of 262.1 h registered indoors while cows visited the dairy milking barn, and 400.4 h registered outdoors while cows remained at pasture. The 133 label files are a list of timestamps indicating the start and end of identified animal behaviors and other annotation remarks. Labels of animal behavior categories include grazing and rumination in standing and lying-down positions, among others. Other annotation labels indicate that the animal is in the barn and the time of installation and removal of the recording systems. The JM label files specify two types of information: (*i*) a list of timestamps indicating the start and end, and type of JMs; (*ii*) a list of timestamps with the middle location and type of JMs.
2. JMs: It consists of a ZIP file containing 2 WAV audio files and label files of JMs. The WAV files correspond to a grazing and rumination bout extracted from channel 1 of the ‘D3RS4ID2909P3.mp3’ file, lasting 54.6 and 30.0 min, respectively. Each WAV file has three associated label files in each format (TXT and CSV file extension):
  - A file generated by the experts indicating a list of timestamps (start and end) and a label with the type of JMs.
  - A label file indicating a list of the middle location (single mark) and the type of JMs. This file was created using a Python script that computes the middle locations as the average of the starts and ends specified by the experts.
  - A label file generated with a Python script indicating a list of automatically adjusted timestamps (start and end) and the type of JMs labeled by the experts.
  The former label files are also provided for direct usage with the “D3RS4ID2909P3.mp3” file.
3. Additional information:
  - The ‘BehaviorLabelsDescription.pdf’ file provides a comprehensive description of animal behavior categories, including the registered annotations and the criteria used by the experts to determine the start and end of each behavior.
  - The ‘JMDescription.pdf’ file explains the marks and characteristics used to distinguish the different ingestive and rumination JMs produced during grazing and rumination activities, respectively.
  - The ‘MP3AudioInformation.xlsx’ file provides three worksheets with detailed information on the audio recordings. Information consists of the corresponding trials of the Latin-square design (day, cow and recording system), date, audio duration, sound quality, registered animal behaviors, audio channels, and companion comments.

**Figure 2.**
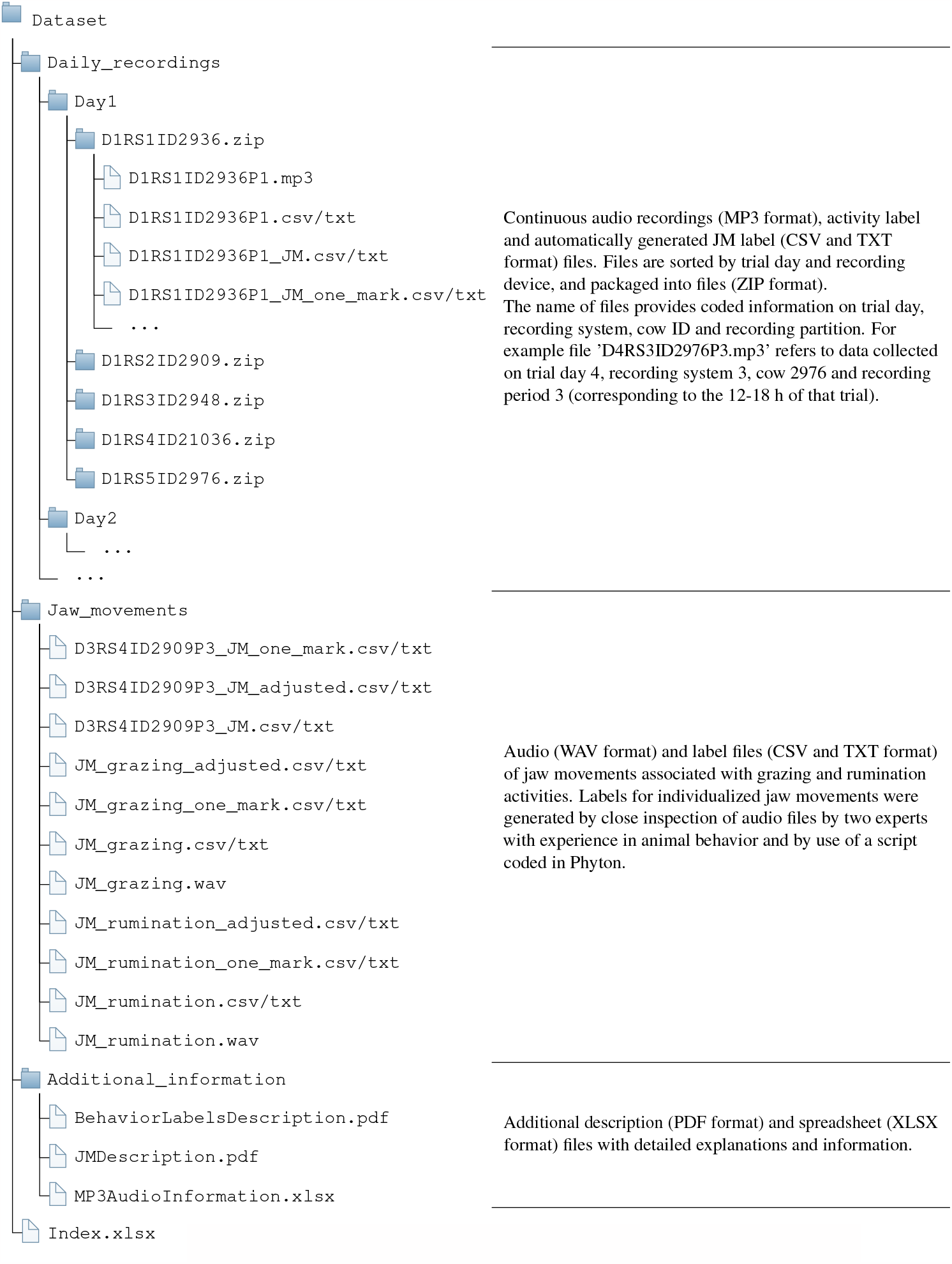
Internal dataset organization in bundled files and naming.

## Technical Validation

The interruptions of regular JMs performed rhythmically in grazing and rumination activities can be used to delimit their bouts^12^. In this study, interruptions of consecutive JMs greater than 90 s were considered to delimit the grazing and rumination bouts. The duration of the grazing and rumination bouts is shown in Fig. 3. Small interruptions between two consecutive grazing bouts could be associated with an animal distraction or animal walking to a distant feeding patch. The great sensitivity to interruptions of regular JMs generates multiple short grazing bouts that can be aggregated into longer grazing meals, making it useful to estimate minute to hourly grazing time budgets. Thus, about 40% of the grazing bouts last less than 25 min (see Fig. 3), while a typical grazing meal lasts more than 1 h^12^. About 85% of the rumination bouts lasts less than 75 min (Fig. 3). The waveform and spectrogram of audio signals during grazing and rumination are shown in Fig. 4a and Fig. 4b, respectively. The bottom panel of Fig. 4b shows a zoom-in of the waveform region produced during the pause required for swallowing and regurgitating the feed cud between two consecutive chewing periods^6,40^. A more detailed explanation of grazing and rumination activities is provided in the file ‘BehaviorLabelsDescription.pdf’.

**Figure 3.**
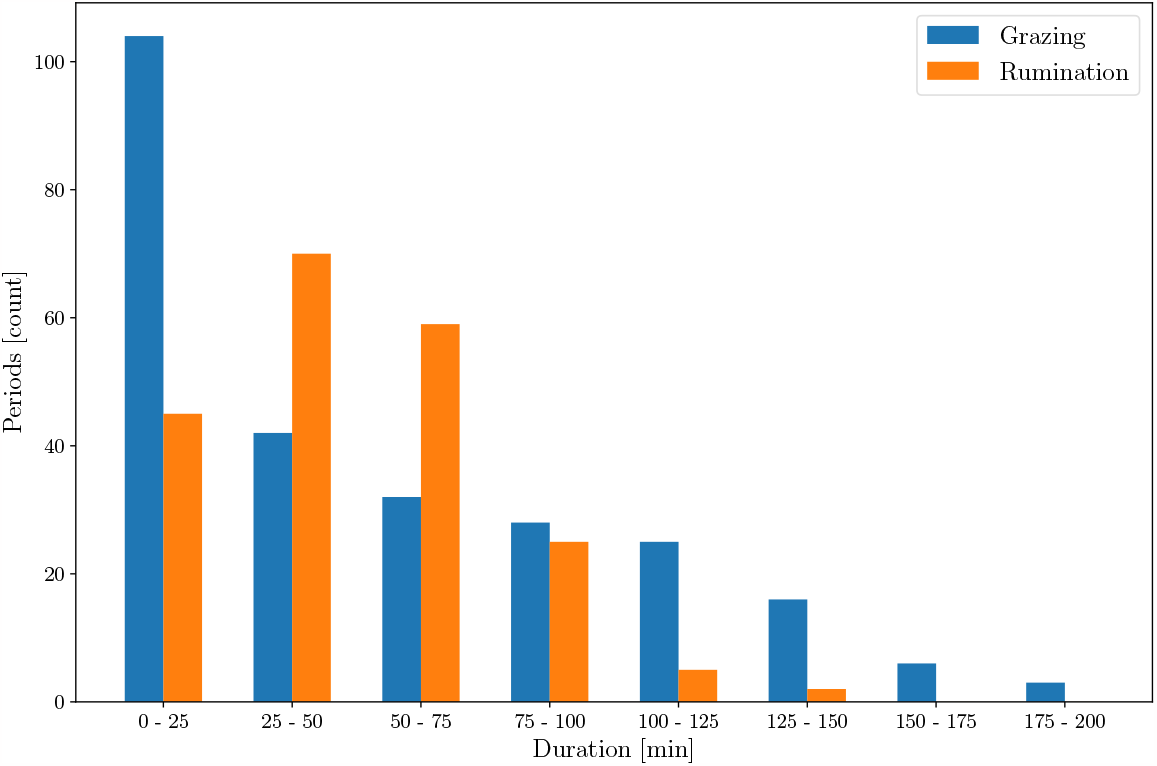
Histogram showing the frequency distribution of the duration of grazing and rumination bouts grouped in 25 min intervals. A total of 257 grazing bouts and 206 rumination bouts are present in the dataset.

**Figure 4.**
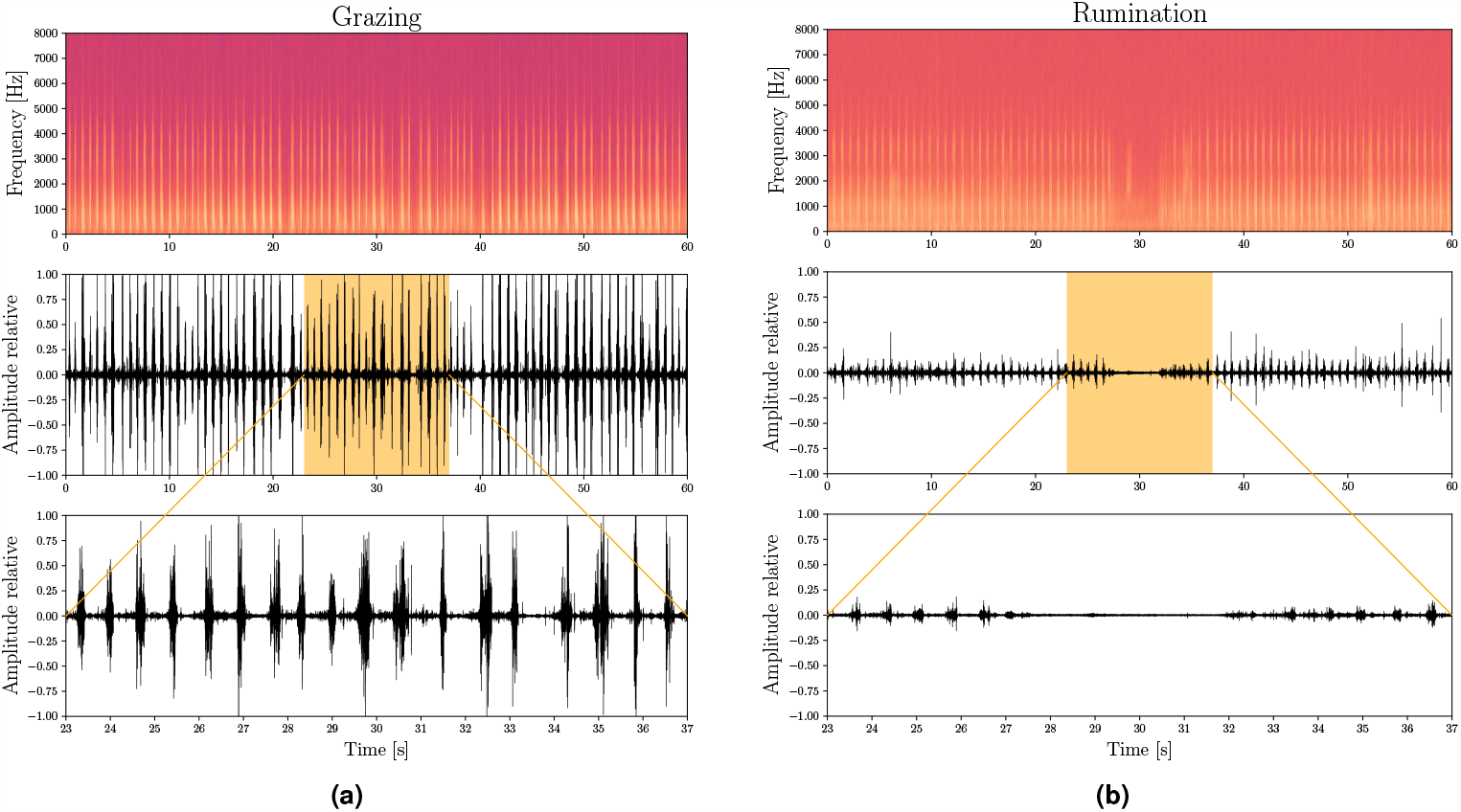
Spectrogram and waveform (with zoom) of foraging audio signals associated with (a) grazing and (b) rumination activities.

The 6,227 JMs labeled by the experts correspond to 2,006 chews during rumination (32.2%), 1,136 chews during grazing (18.2%), 578 bites (9.3%), 2,507 chew-bites (40.3%) and 6 possible non-labeled JMs (<0.1%). This indicates a ratio of chew actions to bite actions performed during grazing of 1.18 (see Equation 1), supporting previously reported results^44^. The number of chews (*N*_*C*_), bites (*N*_*B*_) and chew-bites (*N*_*CB*_) produced in a grazing bout can be used to determine the chew-per-bite ratio (*R*_*C*:*B*_) as:

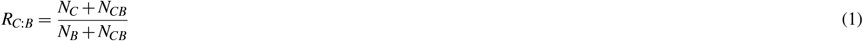

Examples of the waveforms and the average spectral characteristics of the different types of JMs are shown in Fig. 5. A more detailed explanation of the JMs is provided in the ‘JMDescription.pdf’.

**Figure 5.**
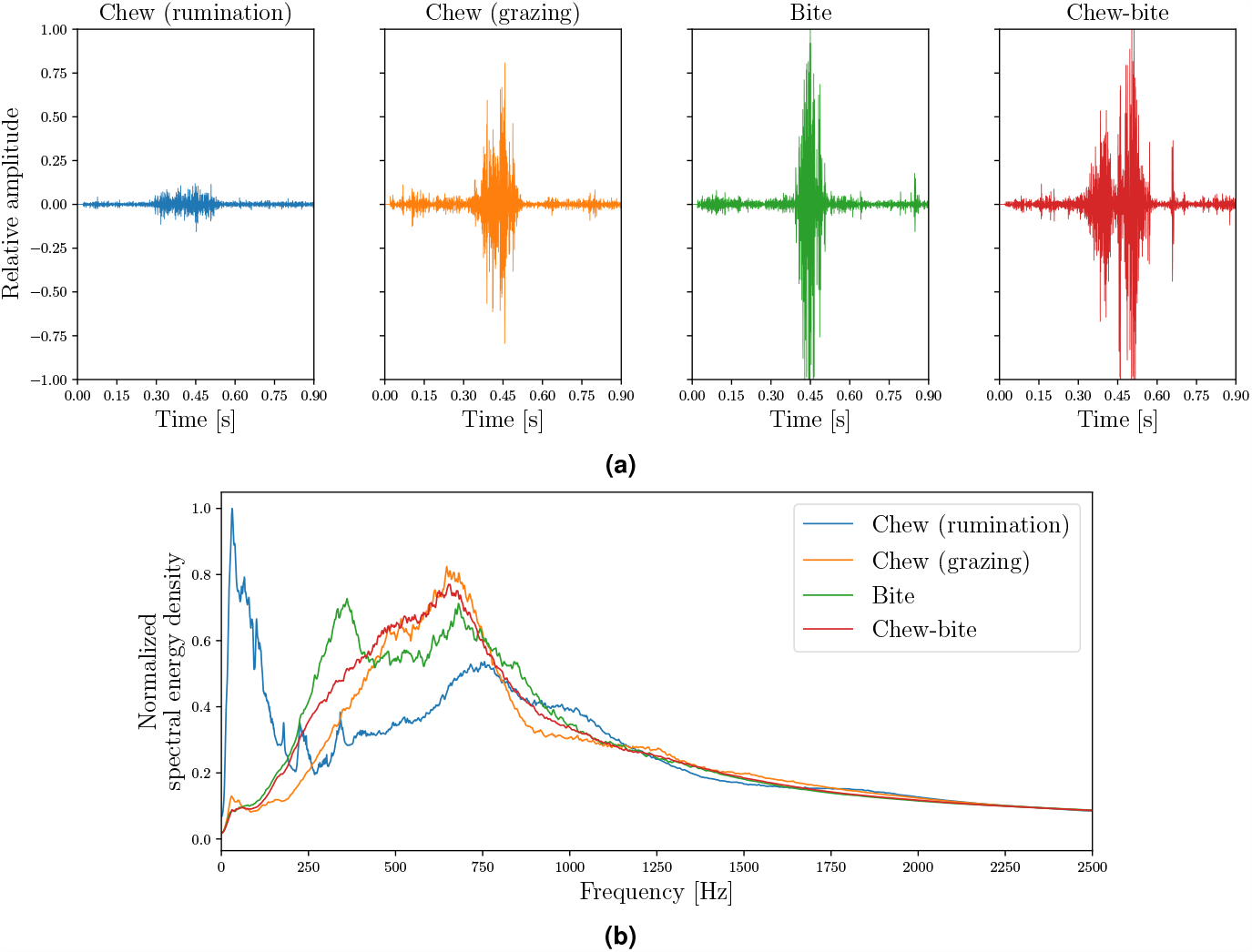
Typical waveform (a) and average spectrum (b) for the different types of JMs: chew produced during rumination and chew, bite and chew-bite produced during grazing. Energy spectra were averaged over all JMs and normalized to the maximum value.

To evaluate the sound quality of the audio recordings obtained from the continuous monitoring of dairy cows, only the active grazing and rumination bouts were examined. Initially, the experts conducted a subjective analysis by listening to random segments of each grazing and rumination bout and confirmed that the corresponding activities were aurally discriminated from the background noise. This statement was further confirmed through a quantitative analysis of these bouts using the JMs’ timestamps automatically generated with the JMs classifier algorithm. For each audio recording, two quality indicators of JMs were individually calculated for grazing and rumination using previously established parameters^30^.

The first parameter, the JM modulation index (MI) is useful to locate the JMs. The MI is a measure based on the difference between the audio signal intensity produced during the JMs and the background noise. Given that the JMs are performed rhythmically every ∼ 1 s during grazing and rumination, the MI was computed as:

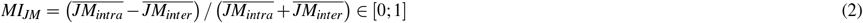

where 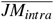 and 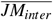 are the mean audio signal intensity produced during JMs and mean audio signal intensity produced in the short-pauses between consecutive JMs respectively, and defined as:

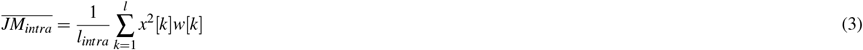

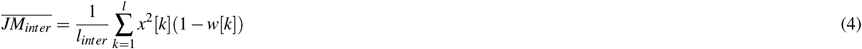

where *x*[*k*] is the audio signal, *l* is the length in samples of the audio signal, *l*_*intra*_ and *l*_*inter*_ are the total number of samples with and without JMs, respectively, and *w*[*k*] is a logical function indicating the presence of a JM in the *k*-th sample.

The second parameter is the signal-to-noise ratio (SNR). This parameter indicates the extent to which the background noise affects the sound produced during JMs, thus helping to differentiate between JMs associated with chews, bites and chew-bites. To compute the SNR, the sound produced during JMs must be isolated from the background noise. A multiband spectral subtraction algorithm assuming uncorrelated additive noise in the audio recordings was used to estimate a noise-free signal *ŝ*[*k*] and a noisy signal 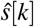[*k*]^45^. The SNR is computed as follows:

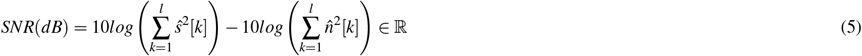

Examples of audio recordings with high- and low-quality sound are available at Gitlab (https://gitlab.com/luciano.mrau/acoustic_dairy_cow_dataset/-/tree/master/data/sound_quality). Their waveforms are presented in Fig. 6a and 6b. The higher the *MI*_*JM*_ and SNR values, the better the audio recording quality. The frequency distribution of the estimated values of *MI*_*JM*_ and SNR for both rumination and grazing computed over the 133 audio recordings of continuous monitoring are shown in Fig. 7. Fig 7a. shows a considerable variation in the *MI*_*JM*_ values of rumination and grazing. The *MI*_*JM*_ values of rumination tend to be smaller than the *MI*_*JM*_ values of grazing. This indicates that the JMs produced in rumination (exclusively chews) are more difficult to distinguish from the background noise. This is partly due to the lower intensity of the rumination JMs compared to the ingestive JMs, as shown in Fig. 4 and Fig. 5a. We hypothesize that the lower intensity in rumination is because of the high moisture content of the ingested matter^30,40^. Fig 7b shows that the ingestive JMs produced during active grazing are less affected by background noise than the rumination JMs produced during rumination. This could be due to the difference in the energy spectral density of the JMs produced in grazing and rumination compared to that of the background noise^46^.

**Figure 6.**
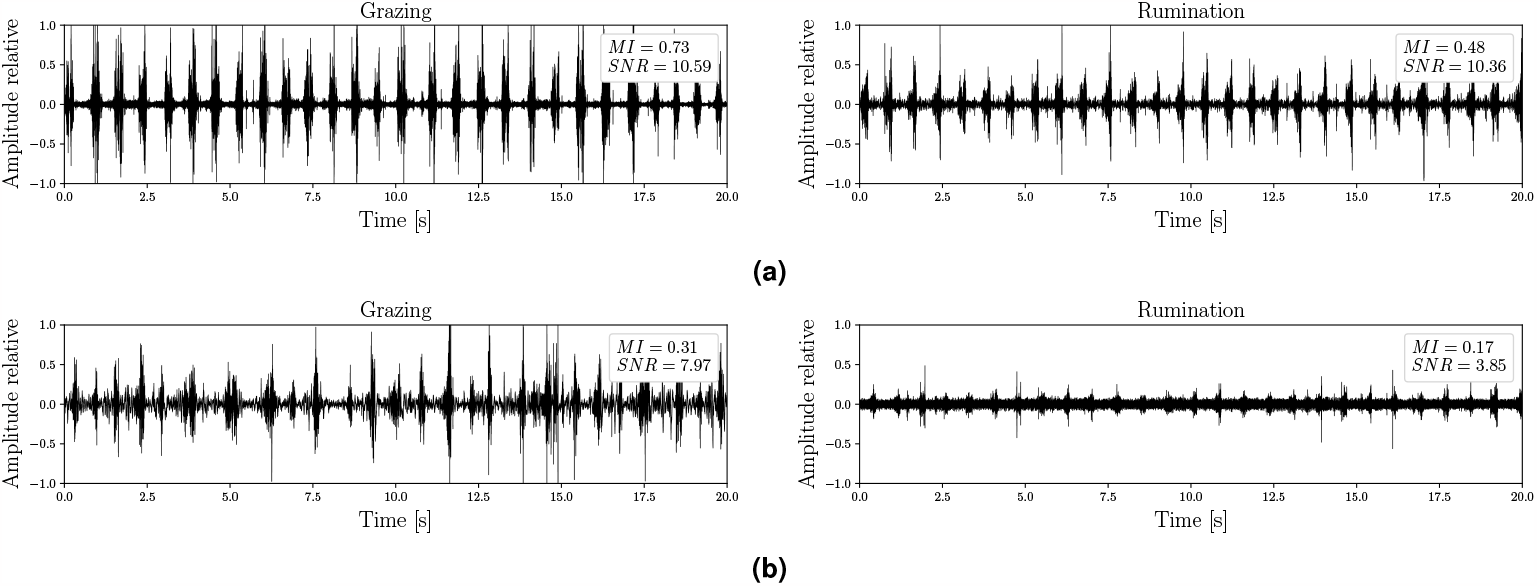
Waveforms of segments of audio recordings with (a) high- and (b) low-quality sound.

**Figure 7.**
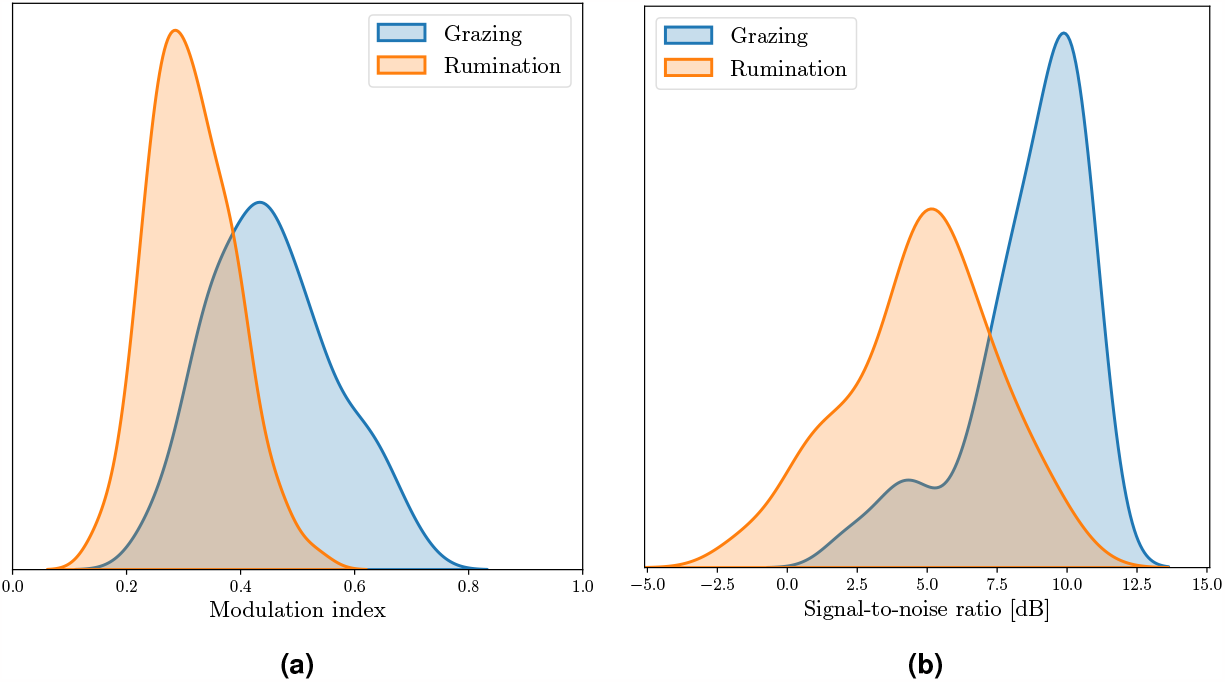
Frequency distribution of the audio recording quality in terms of (a) the modulation index and (b) the signal-to-noise ratio.

Quantitative differences between the two channels of the audio recordings have been measured in terms of the *MI*_*JM*_ and SNR values. Table 4 presents the *MI*_*JM*_ values computed for grazing and rumination in each daily recording. The slash-separated values represent the *MI*_*JM*_ for grazing and rumination. The less the difference in the *MI*_*JM*_ values of channels 1 and 2 of a determined daily recording, the greater the similarity in the signals. Values in blue and orange represent the channels corresponding to the inward- and outward-facing microphone designated by the experts, respectively. Table 5 presents the SNR values in an analogous way to table 4. In particular, the small *MI*_*JM*_ and SNR values of the channel corresponding to the inward-facing microphone from the recordings of day 1 - cow 5, day 3 - cow 3 and day 3 - cow 5 are associated with poor sound quality.

**Table 4.**
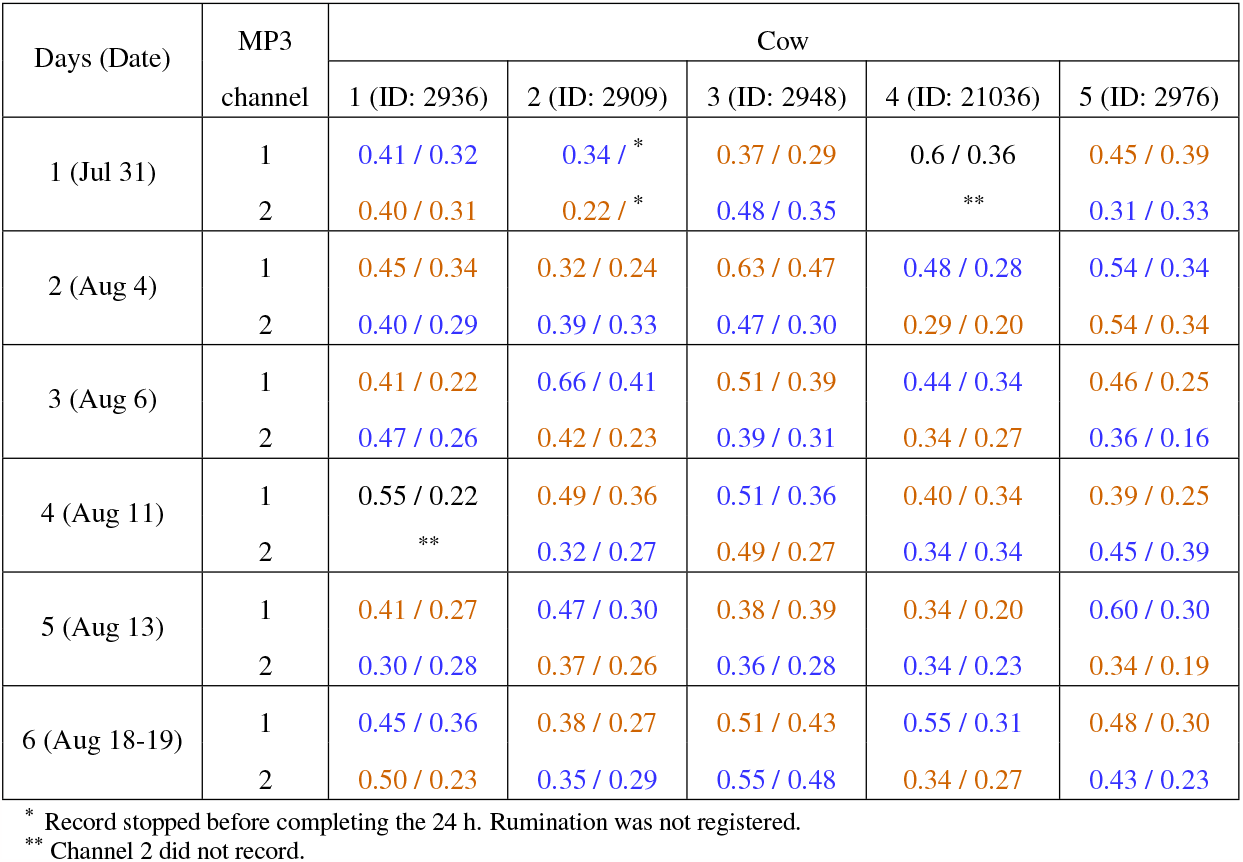
*MI*_*JM*_ values computed in the trials for the two channels of the MP3 files. Separate slash values represent the *MI*_*JM*_ for grazing and rumination. Blue and red color text corresponds to the inward- and outward-facing microphone assignment made by the experts, respectively.

**Table 5.**
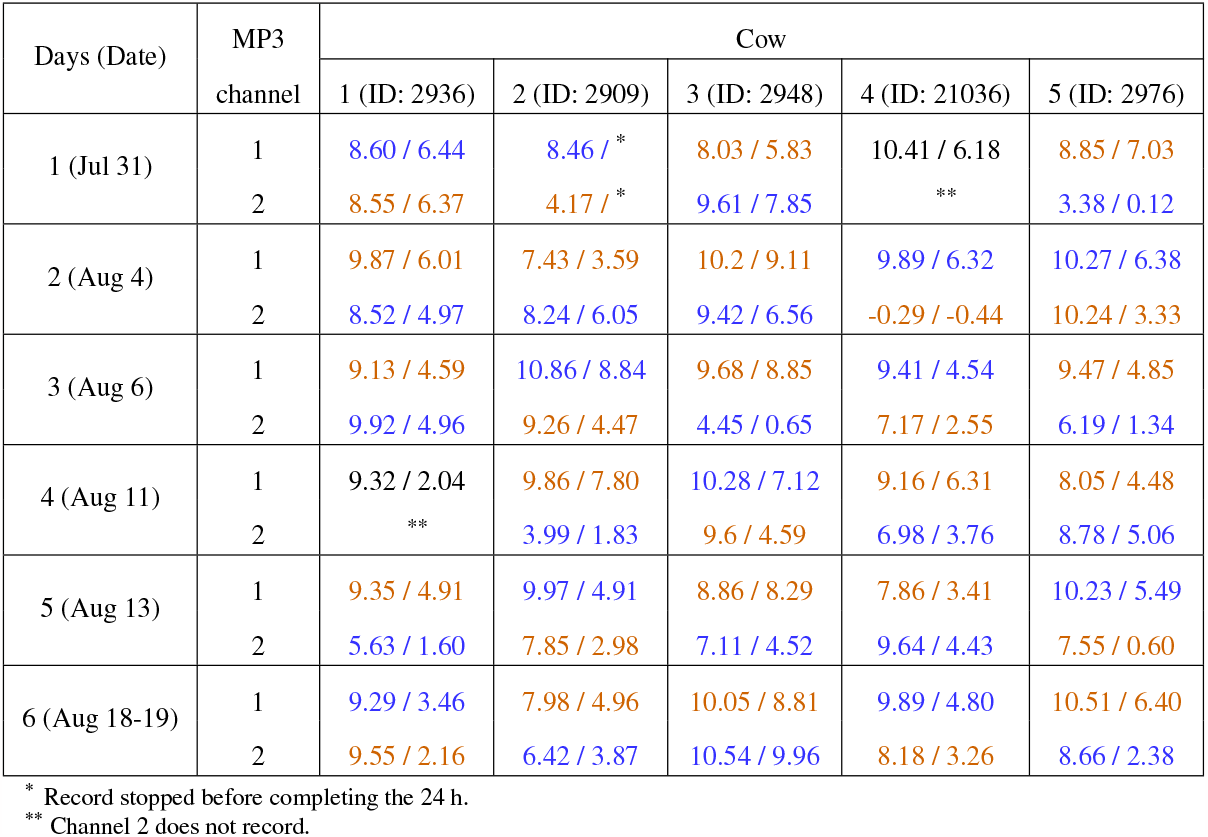
SNR values in dB computed in the trials for the two channels of the MP3 files. Separate slash values represent the SNR for grazing and rumination. Blue and red color text corresponds to the inward- and outward-facing microphone assignment made by the experts, respectively.

| Days (Date) | MP3<br>channel | Cow |  |  |  |  |
| --- | --- | --- | --- | --- | --- | --- |
|  |  | 1 (ID: 2936) | 2 (ID: 2909) | 3 (ID: 2948) | 4 (ID: 21036) | 5 (ID: 2976) |
| 1 (Jul 31) | 1 | 8.60 / 6.44 | 8.46 / * | 8.03 / 5.83 | 10.41 / 6.18 | 8.85 / 7.03 |
|  | 2 | 8.55 / 6.37 | 4.17 / * | 9.61 / 7.85 | ** | 3.38 / 0.12 |
| 2 (Aug 4) | 1 | 9.87 / 6.01 | 7.43 / 3.59 | 10.2 / 9.11 | 9.89 / 6.32 | 10.27 / 6.38 |
|  | 2 | 8.52 / 4.97 | 8.24 / 6.05 | 9.42 / 6.56 | -0.29 / -0.44 | 10.24 / 3.33 |
| 3 (Aug 6) | 1 | 9.13 / 4.59 | 10.86 / 8.84 | 9.68 / 8.85 | 9.41 / 4.54 | 9.47 / 4.85 |
|  | 2 | 9.92 / 4.96 | 9.26 / 4.47 | 4.45 / 0.65 | 7.17 / 2.55 | 6.19 / 1.34 |
| 4 (Aug 11) | 1 | 9.32 / 2.04 | 9.86 / 7.80 | 10.28 / 7.12 | 9.16 / 6.31 | 8.05 / 4.48 |
|  | 2 | ** | 3.99 / 1.83 | 9.6 / 4.59 | 6.98 / 3.76 | 8.78 / 5.06 |
| 5 (Aug 13) | 1 | 9.35 / 4.91 | 9.97 / 4.91 | 8.86 / 8.29 | 7.86 / 3.41 | 10.23 / 5.49 |
|  | 2 | 5.63 / 1.60 | 7.85 / 2.98 | 7.11 / 4.52 | 9.64 / 4.43 | 7.55 / 0.60 |
| 6 (Aug 18-19) | 1 | 9.29 / 3.46 | 7.98 / 4.96 | 10.05 / 8.81 | 9.89 / 4.80 | 10.51 / 6.40 |
|  | 2 | 9.55 / 2.16 | 6.42 / 3.87 | 10.54 / 9.96 | 8.18 / 3.26 | 8.66 / 2.38 |

\* Record stopped before completing the 24 h.
\*\* Channel 2 does not record.

## Usage Notes

Audio editing software, such as Audacity^41^ or Sonic Visualiser^47^, can be used to work with this dataset. The MP3 and WAV files, along with their corresponding label files, can be imported. The multiple label files associated with each audio file (delimited by the experts, delimited automatically or with one central mark) can also be imported simultaneously for comparison or other specific user interests.

The ‘MP3AudioInformation.xlsx’ is a spreadsheet file that provides specific information on the audio recordings obtained from the continuous monitoring of dairy cows. The sheet called “Audiofile properties” describes the Latin-square design for this experiment, which could be useful to analyze variations related to animals, experimental days or recording systems. Additionally, the correspondence between the direction of the microphones (inwards/outwards) and the channels in the audio recordings elaborated by the experts is also indicated. It should be noted that some errors may have occurred in the channel assignment due to the diverse sound quality detected across audio recordings. Any observations or particularities presented in the audio recordings are also mentioned. The sheet named “Cattle activities” specifies the kind of animal behavior categories and annotations presented in the audio recordings. This enables users to filter activities of interest.

Audio recording qualities vary greatly due to differences in microphones and recording channels. We hypothesize that these variations were caused by differences in microphone response, microphone setup at the onset of recordings, and microphone movement during recordings. The sheet named “Audio quality” shows the values of the quality parameters for the audio recordings, using a background color scale from green to red to indicate high- and low-quality sound, respectively. This enables users to choose the optimal audio recordings or apply signal enhancement techniques, among other options. We recommend listening to the audio recordings in stereo or mono, depending on their preferred comfort and result, as this can vary from user to user due to differences in hearing capacity and audio signal intensity. We suggest listening in stereo for audio recordings with high-quality sound and listening only to the channel corresponding to the microphone facing inward for those with low-quality sound, as indicated in the ‘AudioDescription.xlsx’ file.

The information on the JMs labeled by experts can be used as a standalone dataset for JMs analysis and for developing new automatic algorithms for detecting and classifying JMs. We encourage users to utilize the provided JM labels generated by experts as an audiovisual guide and reference to verify and correct the automatically generated JM labels in all audio recordings.

The data described in this article are released under a Creative Commons Attribution-NonCommercial-ShareAlike 4.0 International (CC BY-NC-SA 4.0) license, indicating that it may be used for non-commercial purposes. We encourage users to cite this article when using the data for proper attribution.

## Code availability

The code for automatically adjusting the timesteps of JM labels, computing the JM labels of the audio recordings and for technical validation is available at Gitlab (https://gitlab.com/luciano.mrau/acoustic_dairy_cow_dataset). All code was written in Python 3.8.10 and distributed under the MIT license. Small changes should be made to the scripts by specifying the path of the audio files of the execution environment.

## Acknowledgements

The authors would like to express their gratitude to the staff of the KBS Robotic Dairy Farm for their invaluable support and assistance during the completion of this study. The operation of this farm and the research was possible through funding provided by the USDA-NIFA MICL0222 and MICL0406 projects, and support from Michigan State University AgBioResearch. This work has been funded by various organizations including Universidad Nacional del Litoral, Argentina, with projects CAID 50620190100080LI and 50620190100151LI, Universidad Nacional de Rosario, Argentina, with projects 80020180300053UR, 2016-AGR266 and 2013-AGR216, Consejo Nacional de Investigaciones Científicas y Técnicas (CONICET), Argentina, with project 2017-PUE-sinc(i). Support for weather data was provided by the KBS LTER Program (NSF Award DEB 2224712).

## Author contributions statement

Individual authorships and contributionships are described using the terms described by the Contributor Roles Taxonomy (CRediT) author statement^48^. L.S.M.R: Conceptualization, Methodology, Software, Validation, Formal analysis, Investigation, Data Curation, Writing - Original Draft, Writing - Review & Editing, Visualization. J.O.C: Investigation, Writing - Original Draft, Writing - Review & Editing, Visualization. M.F: Validation, Data Curation, Writing - Review & Editing. S.A.U: Methodology, Design, Formal analysis, Investigation, Data Curation, Writing - Review & Editing. A.M.P: Data Curation, Writing - Review & Editing. L.D.V: Methodology, Writing - Review & Editing. L.L.G: Funding acquisition, Investigation, Methodology, Project administration, Supervision. H.L.R: Funding acquisition, Investigation, Methodology, Project administration, Writing – review & editing, Supervision. J.R.G: Funding acquisition, Investigation, Methodology, Design, Data Curation, Writing - Review & Editing.

## Competing interests

The authors declare no competing financial interests.

## Notes

https://gitlab.com/luciano.mrau/acoustic_dairy_cow_dataset

### Competing Interest Statement

The authors have declared no competing interest.

## References

1. Slob, N., Catal, C. & Kassahun, A. Application of machine learning to improve dairy farm management: a systematic literature review. Prev. Vet. Med. 187, 105237 (2021).

2. Lovarelli, D., Bacenetti, J. & Guarino, M. A review on dairy cattle farming: is precision livestock farming the compromise for an environmental, economic and social sustainable production? J. Clean. Prod. 262, 121409 (2020).

3. Michie, C. et al. The internet of things enhancing animal welfare and farm operational efficiency. J. Dairy Res. 87, 20–27 (2020).

4. Tzanidakis, C., Tzamaloukas, O., Simitzis, P. & Panagakis, P. Precision livestock farming applications (plf) for grazing animals. Agriculture 13 (2023).

5. Banhazi, T. M. et al. Precision livestock farming: an international review of scientific and commercial aspects. Int. J. Agric. Biol. Eng. 5, 1–9 (2012).

6. Hodgson, J. & Illius, A. W. The Ecology and Management of Grazing Systems (Wallingford (United Kingdom) CAB International, 1998).

7. Garcia, R., Aguilar, J., Toro, M., Pinto, A. & Rodriguez, P. A systematic literature review on the use of machine learning in precision livestock farming. Comput. Electron. Agric. 179, 105826 (2020).

8. Aquilani, C., Confessore, A., Bozzi, R., Sirtori, F. & Pugliese, C. Review: precision livestock farming technologies in pasture-based livestock systems. Animal 16, 100429 (2022).

9. Mahmud, M. S., Zahid, A., Das, A. K., Muzammil, M. & Khan, M. U. A systematic literature review on deep learning applications for precision cattle farming. Comput. Electron. Agric. 187, 106313 (2021).

10. Riaboff, L. et al. Predicting livestock behaviour using accelerometers: a systematic review of processing techniques for ruminant behaviour prediction from raw accelerometer data. Comput. Electron. Agric. 192, 106610 (2022).

11. Lovarelli, D. et al. Development of a new wearable 3d sensor node and innovative open classification system for dairy cows’ behavior. Animals 12, 1447 (2022).

12. Andriamandroso, A., Bindelle, J., Mercatoris, B. & Lebeau, F. A review on the use of sensors to monitor cattle jaw movements and behavior when grazing. Biotechnol. Agron. Soc. Environ. 20 (2016).

13. Ferrero, M. et al. A full end-to-end deep approach for detecting and classifying jaw movements from acoustic signals in grazing cattle. Eng. Appl. Artif. Intell. 121, 106016 (2023).

14. Li, G., Xiong, Y., Du, Q., Shi, Z. & Gates, R. S. Classifying ingestive behavior of dairy cows via automatic sound recognition. Sensors 21, 5231 (2021).

15. Duan, G. et al. Short-term feeding behaviour sound classification method for sheep using lstm networks. Int. J. Agric. Biol. Eng. 14, 43–54 (2021).

16. Wang, K., Wu, P., Cui, H., Xuan, C. & Su, H. Identification and classification for sheep foraging behavior based on acoustic signal and deep learning. Comput. Electron. Agric. 187, 106275 (2021).

17. Ungar, E. D. & Rutter, S. M. Classifying cattle jaw movements: comparing iger behaviour recorder and acoustic techniques. Appl. Anim. Behav. Sci. 98, 11–27 (2006).

18. Schirmann, K., von Keyserlingk, M., Weary, D., Veira, D. & Heuwieser, W. Technical note: validation of a system for monitoring rumination in dairy cows. J. Dairy Sci. 92, 6052–6055 (2009).

19. Goldhawk, C., Schwartzkopf-Genswein, K. & Beauchemin, K. A. Technical Note: Validation of rumination collars for beef cattle. J. Animal Sci. 91, 2858–2862 (2013).

20. Martínez Rau, L., Chelotti, J. O., Vanrell, S. R. & Giovanini, L. L. Developments on real-time monitoring of grazing cattle feeding behavior using sound. In 2020 IEEE International Conference on Industrial Technology (ICIT), 771–776 (2020).

21. Laca, E. A., WallisDeVries, M. F. et al. Acoustic measurement of intake and grazing behaviour of cattle. Grass Forage Sci. 55, 97–104 (2000).

22. Galli, J. et al. Monitoring and assessment of ingestive chewing sounds for prediction of herbage intake rate in grazing cattle. Animal 12, 973–982 (2018).

23. Ritter, C., Mills, K. E., Weary, D. M. & von Keyserlingk, M. A. Perspectives of western canadian dairy farmers on the future of farming. J. Dairy Sci. 103, 10273–10282 (2020).

24. Cockburn, M. Review: application and prospective discussion of machine learning for the management of dairy farms. Animals 10 (2020).

25. Vanrell, S. R. et al. Audio recordings dataset of grazing jaw movements in dairy cattle. Data Brief 30, 105623 (2020).

26. Jung, D.-H. et al. Deep learning-based cattle vocal classification model and real-time livestock monitoring system with noise filtering. Animals 11 (2021).

27. Pandeya, Y. R., Bhattarai, B. & Lee, J. Visual object detector for cow sound event detection. IEEE Access 8, 162625–162633 (2020).

28. Deniz, N. N. et al. Embedded system for real-time monitoring of foraging behavior of grazing cattle using acoustic signals. Comput. Electron. Agric. 138, 167–174 (2017).

29. Martinez-Rau, L. S., Weißbrich, M. & Payá-Vayá, G. A 4µw low-power audio processor system for real-time jaw movements recognition in grazing cattle. J. Signal Process. Syst. 95, 407–424 (2023).

30. Martinez-Rau, L. S. et al. A robust computational approach for jaw movement detection and classification in grazing cattle using acoustic signals. Comput. Electron. Agric. 192, 106569 (2022).

31. Chelotti, J. O. et al. An online method for estimating grazing and rumination bouts using acoustic signals in grazing cattle. Comput. Electron. Agric. 173, 105443 (2020).

32. Chelotti, J. O. et al. Using segment-based features of jaw movements to recognise foraging activities in grazing cattle. Biosyst. Eng. 229, 69–84 (2023).

33. Martinez-Rau, L. S., Adin, V., Giovanini, L. L., Oelmann, B. & Bader, S. Real-time acoustic monitoring of foraging behavior of grazing cattle using low-power embedded devices. In 2023 IEEE Sensors Applications Symposium (SAS), 01–06 (2023).

34. Bishop, C. M. Pattern Recognition and Machine Learning (Springer Verlag, 2006).

35. Meng, T., Jing, X., Yan, Z. & Pedrycz, W. A survey on machine learning for data fusion. Inf. Fusion 57, 115–129 (2020).

36. Watt, L. et al. Differential rumination, intake, and enteric methane production of dairy cows in a pasture-based automatic milking system. J. Dairy Sci. 98, 7248–7263 (2015).

37. Milone, D. H., Galli, J. R., Cangiano, C. A., Rufiner, H. L. & Laca, E. A. Automatic recognition of ingestive sounds of cattle based on hidden markov models. Comput. Electron. Agric. 87, 51–55 (2012).

38. Chelotti, J. O. et al. A real-time algorithm for acoustic monitoring of ingestive behavior of grazing cattle. Comput. Electron. Agric. 127, 64–75 (2016).

39. Chelotti, J. O., Vanrell, S. R., Galli, J. R., Giovanini, L. L. & Rufiner, H. L. A pattern recognition approach for detecting and classifying jaw movements in grazing cattle. Comput. Electron. Agric. 145, 83–91 (2018).

40. Galli, J. R. et al. Discriminative power of acoustic features for jaw movement classification in cattle and sheep. Bioacoustics 29, 602–616 (2020).

41. Audacity Team, https://www.audacityteam.org/download/ (2023).

42. Martinez-Rau, L. S. et al. Open dataset of acoustic recordings of foraging behavior in dairy cows. figshare https://doi.org/XXXXX (2023).

43. Bosi, M. & Goldberg, R. E. MPEG-1 Audio, 265–313 (Springer US, Boston, MA, 2003).

44. Ungar, E. D. et al. The implications of compound chew–bite jaw movements for bite rate in grazing cattle. Appl. Animal Behav. Sci. 98, 183–195 (2006).

45. Loizou, P. C. Speech Enhancement: Theory and Practice (CRC press, 2013).

46. Oppenheim, A. V., Willsky, A. S., Nawab, S. H. & Ding, J.-J. Signals and Systems, vol. 2 (Prentice hall Upper Saddle River, NJ, 1997).

47. Cannam, C., Landone, C. & Sandler, M. Sonic visualiser: An open source application for viewing, analysing, and annotating music audio files. In Proceedings of the 18th ACM International Conference on Multimedia, 1467–1468 (2010).

48. Brand, A., Allen, L., Altman, M., Hlava, M. & Scott, J. Beyond authorship: attribution, contribution, collaboration, and credit. Learn. Publ. 28, 151–155 (2015).

